# Analysis of MIKC^c^-type MADS-box genes and proteins in the fern *Ceratopteris richardii*

**DOI:** 10.64898/2026.06.19.732661

**Authors:** Derry Carr, Kirsty McCready, Andrew R. G. Plackett, Juliet C. Coates

## Abstract

The MIKC^c^ subfamily of MADS-box proteins play a key role in flowering plant reproduction, specifying and patterning the floral organs. MIKC^C^ genes have been identified in non-flowering plants, the number increasing with the amount of whole-genome sequencing information available, but MIKC^c^ functions outside flowering plants are less well understood. In this study, we have cloned and sequenced 14 of the 21 MIKC^c^ genes in the model fern *Ceratopteris richardii* and identified expressed cDNAs/transcripts for a further 6 genes, extending and correcting previous genome-and transcriptome analysis. We have identified that the majority of *Ceratopteris* MIKC^c^ genes are expressed in the reproductive sporophyte, with some genes showing reproductive specificity. Using protein alignments and structural modelling, we have shown that *Ceratopteris* MIKC^C^ proteins are structurally diverse, with over half the proteins possessing extended regions N-terminal to the MADS DNA binding domain, suggesting divergent functions or regulation.

## Introduction

The evolution of seeds approximately 370 million years ago (MYA) in the late Devonian period was a key land plant innovation. The reproductive changes leading to seed formation enabled sexual reproduction independently of water, with subsequent desiccation resistance of the enclosed embryo, dispersal over long distances and the state of dormancy, all of which contributed to the dominance of seed plants (Spermatophyta) in most ecosystems (Finch-Savage and Leubner-Metzger 2006; Harrison and Morris 2018; Linkies et al. 2010; Woudenberg et al. 2024; Sharma et al. 2021). Building on this innovation was the advent of the flowering plants (Angiosperms) (Ma et al. 2025; Magallon et al. 2015; Smith and Beaulieu 2024), with more complex reproductive structures for pollination and fertilisation and seeds with endosperm storage tissue (Baroux and Grossniklaus 2019; Baroux et al. 2002; Sharma et al. 2021). Extant seed plants, particularly flowering plants, are critical for human survival. Globally, more than 40% of calories consumed come from just 3 seed crops: rice, wheat and maize (FAO 2018). Additional key seed crops include other cereals (e.g. rye, oats, millet) and pulses (FAO 2024a; Ritchie et al. 2023). Seed plants, particularly cereals, are also critical as fodder for farmed animals consumed by humans (FAO 2024b).

The molecular mechanisms underpinning the development of flowers and seeds have been well-studied in flowering plants including the genetic model *Arabidopsis thaliana*. One key set of genes required for floral specification and patterning is the MADS (MCM1, AGAMOUS, DEFICIENS, SRF1)-box family (Bowman and Moyroud 2024; Chanderbali et al. 2016; Coen and Meyerowitz 1991; Theissen et al. 2016). Based on their domain structure, MADS-box proteins fall into two major classes, Type I and Type II, with Type II MADS-box proteins further divided into MIKC* and MIKC^c^ (Becker and Theissen 2003; Gramzow and Theissen 2010; Lai et al. 2019). While the MIKC* proteins are proposed to function in the gametophyte of flowering plants (embryo-sac, pollen) and of other land plants (Kwantes et al. 2012), the MIKC^c^ subfamily of proteins have vegetative and reproductive functions in the flowering plant sporophyte (Zhang et al. 2024a), most famously specifying floral organ identity: development of the sepals, petals, stamens, carpels and ovules (Bowman and Moyroud 2024; Theissen et al. 2016). In *Arabidopsis*, the A-class gene *APETALA1* (*AP1*) specifies sepals along with the non-MADS gene *APETALA2* (*AP2*). A-class genes, in conjunction with the B-class genes *APETALLA3* (*AP3*) and *PISTILLATA* (*PI*) specify petals (reviewed in Theissen et al 2016). The C-class gene *AGAMOUS* (*AG*) specifies carpels by itself and stamens with the B-class genes. The *AGAMOUS*-related *SEEDSTICK* (*STK*) and *SHATTERPROOF1* and *-2* (*SHP1* and *-2*) D-class genes specify ovules inside the carpels while the redundant E-class genes (*SEPALLATA 1-4*) are required for A, B, C and D function (reviewed in (Theissen et al. 2016)). The floral organ identity genes show precisely regulated expression in the flower primordium (reviewed in (Jack 2004)) and protein products of the A, B, C, D and E-class genes form specific tetramers upstream of the genes required to produce each floral organ - the floral quartet model (Theissen et al. 2016; Theissen and Saedler 2001). While species-specific variations are found based on precise floral morphology and the organs present (Bowman and Moyroud 2024; Theissen et al. 2016), this underlying model of floral organ specification remains conserved across flowering plants.

MIKC^c^ MADS-box protein structure consists of a conserved DNA-binding MADS domain (Schwarz-Sommer et al. 1990), an intervening I domain conferring DNA-binding strength and specificity (Lai et al. 2021), a K domain mediating dimerization and quartet formation (Lai et al. 2019; Puranik et al. 2014) and a C domain that is not well conserved in sequence between proteins and may be involved in transcriptional activation (Cho et al. 1999; de Folter et al. 2005; Goto et al. 2001). Dimerisation and subsequent higher-order tetramerisation is important for MIKC^c^ protein DNA-binding function to regulate flower patterning (Hugouvieux et al. 2024; Immink et al. 2010; Immink et al. 2009; Jack 2004; Puranik et al. 2014). Studies in yeast (two-, three-and four-hybrid) and *in vitro* showed that *Arabidopsis* MIKC^c^ proteins form a range of specific heterodimers and a smaller repertoire of homodimers (de Folter et al. 2005; Immink et al. 2009; Jack 2004; Lai et al. 2021; Melzer and Theissen 2009; Melzer et al. 2009; Puranik et al. 2014; Tonaco et al. 2006; Riechmann et al. 1996; Honma and Goto 2001) that support the floral quartet model.

Despite a key role in floral organ specification, the MIKC^c^ MADS-box gene family is not unique to flowering plants, being present throughout land plants (Thangavel and Nayar 2018). MIKC^c^ genes most likely arose from a single common ancestor predating land plants (Gramzow et al. 2012). Charophycean algae possess a single MIKC^c^ gene that may function in haploid reproductive cell differentiation (Tanabe et al. 2005). In the non-flowering seed plants, gymnosperms, the range of MIKC^c^ genes present is different to that in flowering plants. For example, A-function genes and *SEPELLATAs* are absent and some gymnosperm-specific clade expansions have occurred (Chen et al. 2017). Despite gymnosperm cones being non-homologous with angiosperm flowers (O’Maoileidigh et al. 2014), gymnosperm B and C class MIKC^c^ MADS-box genes have been implicated in reproductive development where it is suggested that B and C class genes together direct male cone development while C class MIKC^c^ genes direct female cone development, with corresponding shifts in B and C class MIKC^c^ gene expression seen during bisexual cone development (Feng et al. 2024; Wang et al. 2010). As in flowering plants, gymnosperm MIKC^c^ proteins can form both homo- and hetero-tetramers: it has therefore been proposed that the floral quartet mechanism arose before flowering plants, as a result of interaction between pairs of homodimers bound to DNA (Melzer et al. 2010).

Knowledge of MIKC^c^ function in ancestral seedless (spore-bearing) plants is limited compared to the wealth of flowering plant analysis. The liverwort *Marchantia polymorpha* possesses a family of Type II MADS-box genes (Bowman et al. 2017), comprising a single MIKC^c^ gene preferentially expressed in gametophyte gametangia (Dipp-Alvarez et al. 2025) and a single MIKC* gene that forms homodimers (Zobell et al. 2010). In the moss *Physcomitrium patens* the six MIKC^c^ genes show relatively widespread expression in the sporophyte and gametophyte (Barker and Ashton 2013; Hohe et al. 2008; Koshimizu et al. 2018; Quodt et al. 2007; Singer et al. 2007) while knock-down and loss-of-function studies implicate *Physcomitrium* MIKC^c^ genes in the formation of gametophyte reproductive organs (sperm, antheridia and archegonia) and subsequent sporophyte development, but also in gametophyte vegetative development (Koshimizu et al. 2018; Singer et al. 2007). Within seedless vascular plants, lycophyte MIKC^c^ genes show broad expression across sporophyte development (Svensson and Engstrom 2002; Tanabe et al. 2003). In ferns, pre-genomic studies identified and cloned 17 Type II MADS-box genes (13 MIKC^c^ and 4 MIKC*) from *Ceratopteris richardii* (hereafter ‘*Ceratopteris’*) (Hasebe et al. 1998; Kofuji and Ysmaguchi 1997; Kwantes et al. 2012; Munster et al. 1997; Theissen et al. 2000). Expression of several *Ceratopteris* MIKC^c^ genes was detected by northern blot in both the gametophyte and sporophyte (*CRM1*, *CRM3*/*CMADS6*, *CMADS2*, *CMADS3*) (Hasebe et al. 1998; Munster et al. 1997) while *CMADS1* was detected only in sporophyte tissues, showing enriched expression in sporophylls, i.e. sporangium-bearing reproductive fronds (shoot lateral organs analogous to leaves) (Hasebe et al. 1998). Furthermore, *CMADS1* was localised by *in situ* hybridisation to developing sporangia and its encoded protein was proposed to share structural similarity with the class C gene *AGAMOUS* (Hasebe et al. 1998). Overexpression of *CRM3/CMADS6*, reported as having structural similarities to flowering plant B-class MADS-box proteins and implicated in frond development, disrupts sporophyll elongation (Schulz and Theissen 2025), suggesting a possible ancestral role in development of seedless plant reproductive tissues.

A partial *Ceratopteris richardii* genome (38% coverage) was published with an accompanying transcriptome (Marchant et al. 2019). The genome was then updated with a new assembly and additional transcriptome data (Marchant et al. 2022). The updated genome identified 35 MADS box genes, of which 21 are MIKC^c^ genes (Marchant et al. 2022). Two MIKC^c^ genes exhibit alternative splicing to encode proteins with alternate MADS box domains (Marchant et al. 2022). There has been loss of an ‘orphan’ clade of fern-specific MADS-box genes in *Ceratopteris* (Marchant et al. 2022; One Thousand Plant Transcriptomes 2019). The last common ancestor of ferns and seed plants likely had around 3 Type I MADS-box genes and around 5-6 Type II MADS-box genes (Zhang et al. 2024). There has been a large expansion of MIKC^c^ genes in leptosporangiate ferns such as *Ceratopteris*, although the numbers are on average lower than in seed plants (Zhang et al. 2024). However, to our knowledge, no systematic structural characterisation of MIKC^c^ proteins has been carried out to date in the fern family.

In this paper we have cloned, sequenced and characterised the expression patterns of the *C. richardii* MIKC^c^ MADS-box genes. We have provided insights into the likely protein structure of the *Ceratopteris* MIKC^c^ proteins compared to *Arabidopsis*.

## Results

### Identification and cloning of MIKC^c^ MADS box genes in the *Ceratopteris richardii* genome

The most recent *Ceratopteris* genome annotation identified 21 MIKC^c^ MADS box genes, of which 12 had transcripts detected by RNAseq (Marchant et al., 2022; Figure 1), adding 6 *Ceratopteris* MIKC^c^ cDNA sequences to those found in pre-genomic studies in *C. richardii* and *C. pteroides* (Hasebe et al., 1998; Munster et al., 2007, Kofuji and Yamaguchi 1997; Theissen et al., 2000; Figure 1). Overall, using primers designed to amplify predicted full-length cDNAs, we were able to PCR-amplify 16 *Ceratopteris* MIKC^c^ cDNAs from RNA generated from whole gametophyte and/or reproductive sporophyte (sporophyll) tissue (Figure 1; Supplemental Figure 1). Of the 9 MIKC^c^ genes with no previously-detectable transcripts (Marchant et al., 2022) or for which only partial cDNAs had been isolated (*CRM5*/*7*, Munster et al., 1997), we successfully amplified full-length cDNAs for a further 3 genes, namely *CericMADS7*, *CericMADS10* and *CericMADS23*, although *CericMADS23* was not cloned and sequenced (Figure 1; Supplemental Figure 1; Supplemental Table 1). Of the 16 candidate full-length cDNAs we could PCR-amplify, we were able to clone and sequence 14 full-length MIKC^c^ MADS-box genes (Figure 1; Supplemental Table 1). Of these 14 cloned and sequenced genes, 2 genes, *CericMADS10* (homologous to *CRM5* in *C. pteroides* (Munster et al. 1997)) and *CericMADS24* were cloned in full for the first time. The majority of cloned genes were identical in sequence to the published genome ((Marchant et al. 2022); Supplemental table 1). However, cloning and sequencing of *CericMADS10* revealed a 3-nucleotide deletion in the full-length transcript compared to the most recently annotated genome sequence (which assembled two partial transcripts) resulting in a single-amino acid deletion in the predicted protein-coding sequence (Figure 2). This analysis thus experimentally validated the existence of previously-predicted but unverified MIKC^C^ MADS-box genes in *Ceratopteris*.

**Figure 1:**
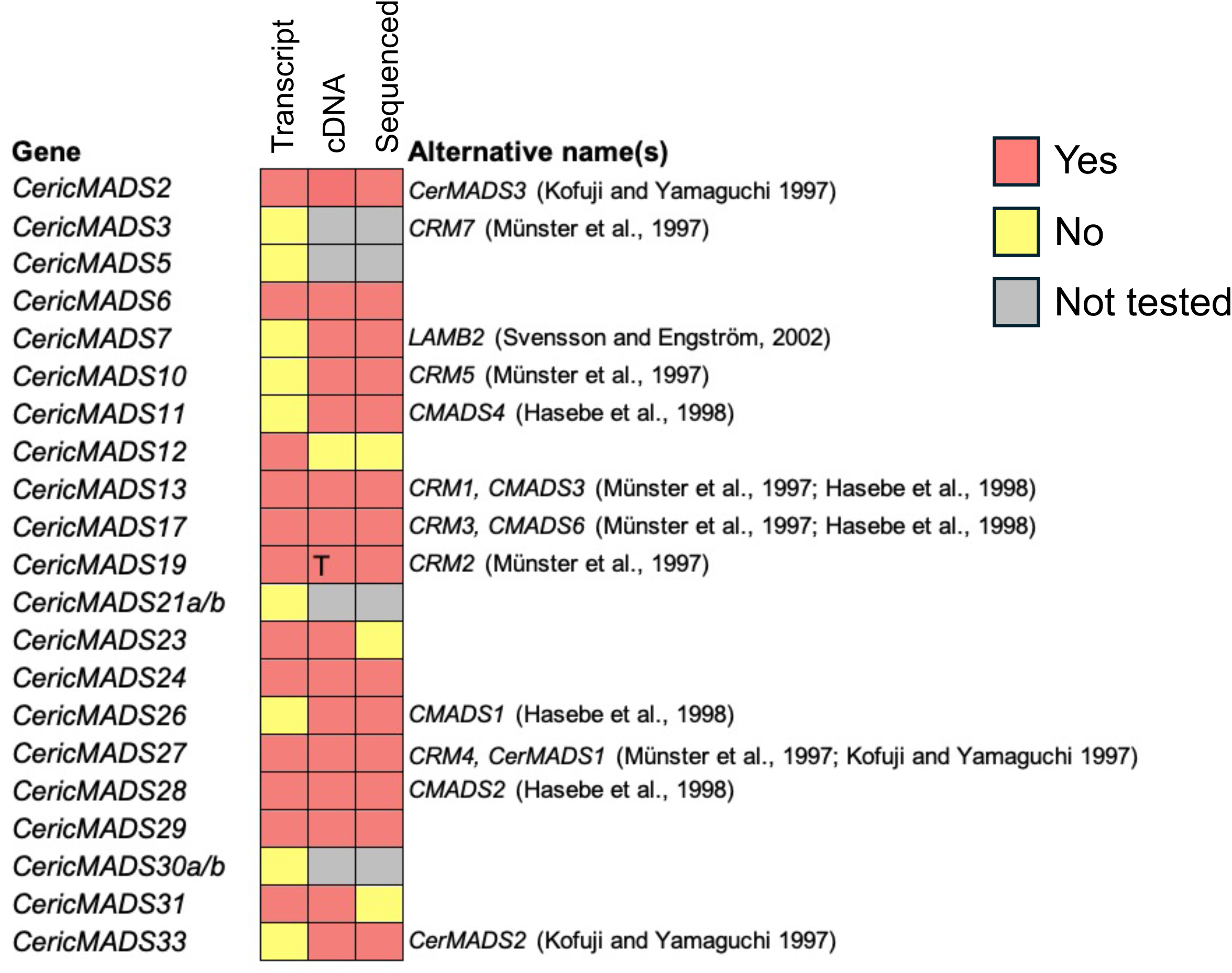
Cloning and identification of *Ceratopteris* MIKC^c^ MADS-box genes and consolidation with previously published data. Gene nomenclature from the most recent genome annotation (Marchant et al., 2022) is shown on the left. From left to right, the heatmap shows the presence or absence of a transcript detected by RNAseq in the most recent transcriptome analysis (Marchant et al,. 2022), presence or absence of a full-length cDNA PCR-amplified from *Ceratopteris* RNA samples in this study, and whether the cDNA was cloned into the pJET cloning vector and sequenced. Red indicates a positive result (transcript, cDNA amplification or cloning/sequencing), yellow indicates a negative result (for cDNA amplification, no signal detected in 3 replicates), and grey indicates genes that were not tested in subsequent steps. ‘T’ indicates that the cDNA amplified was truncated compared to the genome annotation (Marchant *et al*., 2022). Previous alternative names for the genes with relevant references are shown to the right of the heatmap.

**Figure 2.**
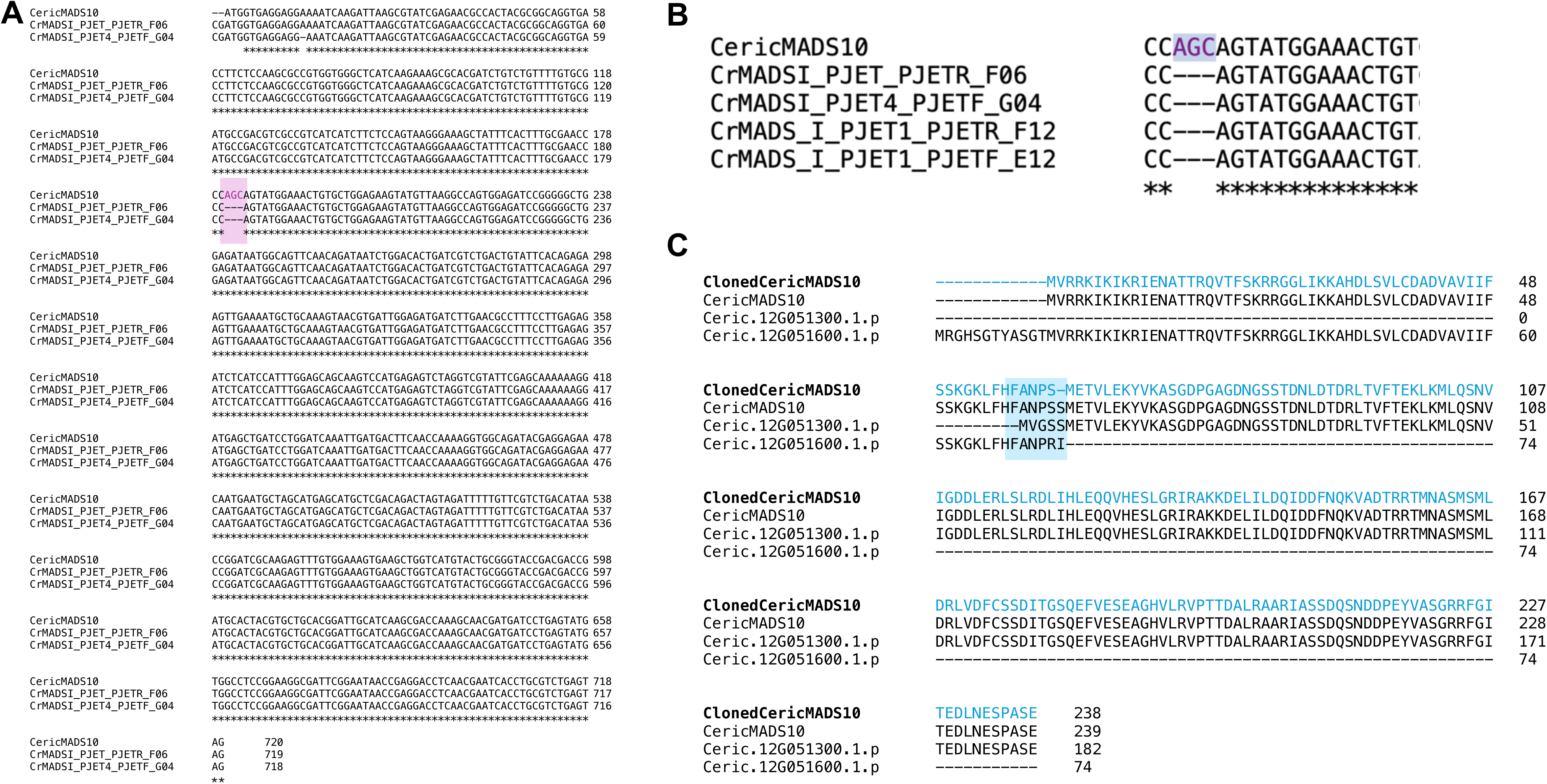
Cloning and sequencing of *CericMADS10*. A. RT-PCR amplification of full-length *CericMADS10* from gametophytic and reproductive tissues. Three biological repeats are shown with a no reverse transcriptase (-RT) control. Arrowhead, 700bp. A selection of these amplified products were cloned into PJET and sequenced. B. 2 sequencing runs (forward and reverse) from PCR-amplified and cloned *CericMADS10* in PJET (CrMADSI_PJET_PJETR_F06 and CrMADSI_PJET4_PJETF_G04) suggested a deletion compared to the published *CericMADS10* sequence (labelled as CericMADS10). The discrepancy/insertion in *CericMADS10* is highlighted in pink with purple text. C. Further sequencing (forward and reverse) and alignment of the deleted *CericMADS10* region from an independent clone in pJET (sequences CrMADS_I_PJET1_PJETR_F12; CrMADS_I_PJET1_PJETF_G12) demonstrated the deletion in the same area compared to the CericMADS10 genome sequence (purple text). D. Protein sequence alignment from translation of the published version of CericMADS10 (Merchant et al., 2022), translation of the two transcripts encoding CericMADS10 fragments from Phytozome (Ceric.12G0513001.1.p and Ceric.12G0516001.1.p) and translation of the cloned CericMADS10 sequence. The area of discrepancy between the translated sequences is highlighted in light blue, and the newly cloned *Ceric*MADS10 sequence (ClonedCericCMADS10) is shown in blue.

### *Ceratopteris* MIKC^c^ genes are expressed in gametophytes and sporophytes, including reproductive tissue

Previous transcriptome analysis has suggested most *Ceratopteris* genes are expressed in both the gametophyte and sporophyte stage (Marchant et al. 2022). Flowering plant MIKC^c^ MADS genes pattern the flower including the ovule, where the female gametophyte and then the seed arises. Seeds are made of both maternal reproductive sporophyte tissue and gametophyte-derived tissue. Thus, we examined whether we could detect expression of full-length *Ceratopteris* MIKC^c^ cDNAs in three stages of reproductive sporophyte tissue (young fiddleheads, unfurling fronds and mature spore-bearing fronds) using RT-PCR (Figure 3A; Supplemental Figure 1).

**Figure 3.**
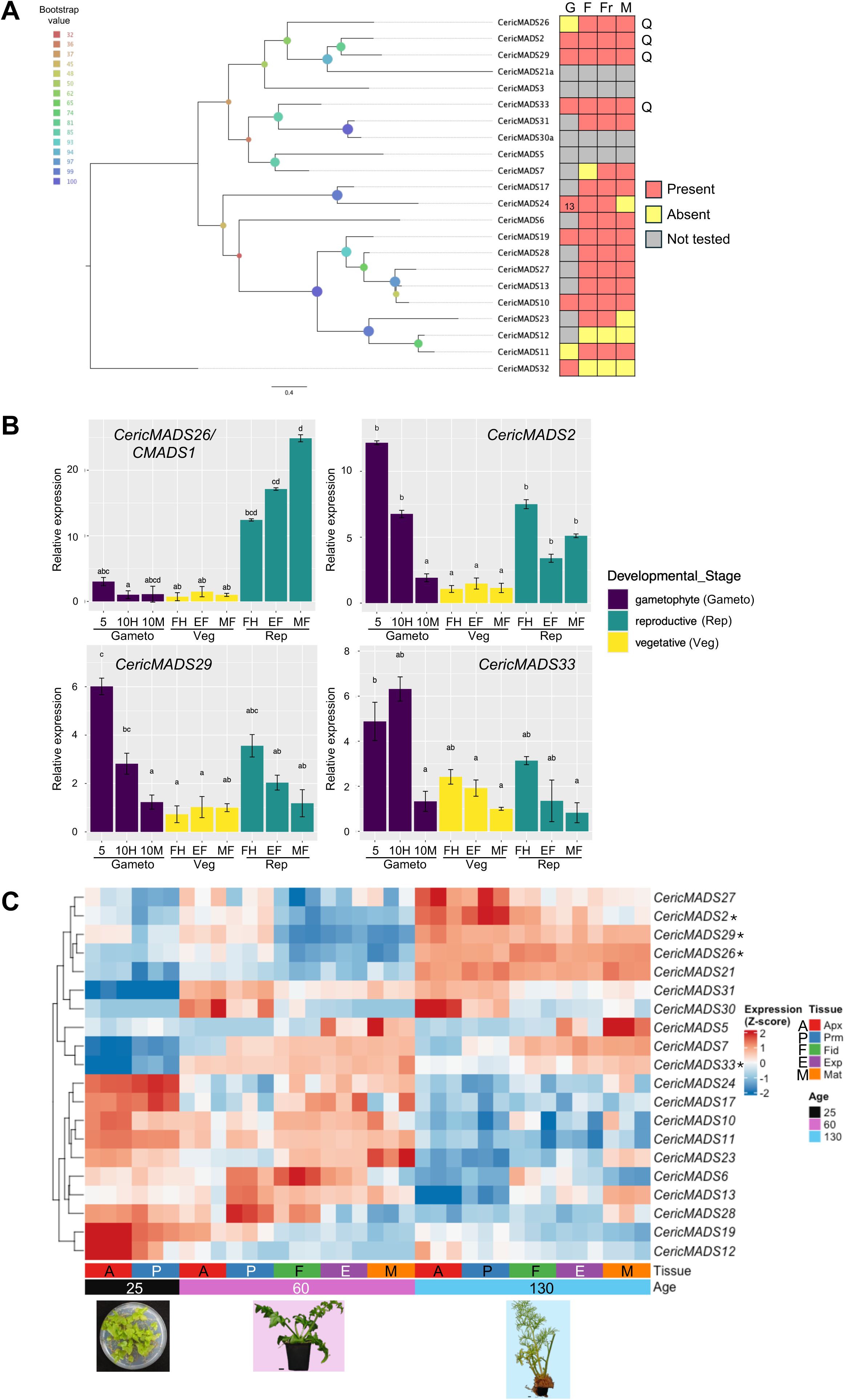
Expression analysis of *Ceratopteris* MIKC^c^ genes. A. Phylogeny reconstruction of *Ceratopteris* MIKC^c^ proteins with the MIKC* protein *Ceric*MADS32 as an outgroup is shown on the left. The bootstrap score at each node is shown in colour according to the legend on the left. The heatmap on the right shows the presence (red) or absence (yellow) of cDNA in gametophyte (6 days and 13 days except *Ceric*MADS24) (G), reproductive fiddlehead (F), unfurling reproductive fronds (Fr) and mature reproductive fronds (M). Grey, not tested. The genes marked with ‘Q’ were selected for qRT-PCR analysis. B. Expression analysis of *Ceric*MADS26/CMADS1 and the related genes *CericMADS2*, C*ericMADS29* and C*ericMADS33* by qRT-PCR. Purple, gametophyte; yellow, vegetative (60d) tissue; green, reproductive (130d) tissue. 5, 5-day; 10H, 10-day hermaphrodite; 10M, 10-day male; FH, fiddlehead; EF, expanding frond; MF, mature frond. Relative expression for each gene was calculated using the “vegetative mature frond” developmental stage as the reference, due to all 4 genes having low expression at this stage. For each gene ANOVA and a post-hoc Tukey statistical test were used to determine significant differences in expression between tissue types: significance is indicated by the letters above each tissue type bar with the same letter above different bars indicating no significant difference between the tissue types. Error bars are the standard error of the delta CT values of the 3 biological repeats performed with each tissue type; each biological repeat is the average of 4 technical repeats. C. Expression profiles of the 20 *Ceratopteris* MIKC^c^ genes extracted from the sporophyte developmental dataset generated by Plackett and Catoni (2025). Tissue samples include 25d sporophyte (black) apices (Apx; red) and frond primordia (Prm; blue), 60d vegetative sporophyte (pink) apices, frond primordia, fiddleheads (Fid; green), expanding fronds (Exp; purple) and mature fronds (Mat; orange), and 130d reproductive sporophyte (blue) apices, frond primordia, fiddleheads, expanding fronds and mature fronds. Representative plant images are shown. Three biological replicates are shown for each tissue. Genes were hierarchically clustered (Euclidean distance, complete linkage). Expression values are row-scaled z-scores (Supplemental Table 4). Combined values for *CericMADS21a/b* and *CericMADS30a/b* (shown as *CericMADS21* and *CericMADS30*) were calculated based on combined exonic size. * Genes also tested by qPCR in figure 3B.

We also tested a representative selection of genes from across the MIKC^c^ family for expression during gametophyte development and tested the expression of a MIKC* gene, *CericMADS32* for comparison (Figure 3A; Supplemental Figure 1). We found that the majority of MIKC^c^ genes tested were expressed throughout reproductive sporophyte development (Figure 3A; Supplemental Figure 1). We did not detect *CericMADS7* in young fiddleheads by RT-PCR, nor did we detect *CericMADS23* and *CericMADS24* in mature fronds (Figure 3A; Supplemental Figure 1). Only *CericMADS12* was undetectable in all reproductive sporophyte stages (Figure 3A; Supplemental Figure 1). In contrast, the MIKC* *CericMADS32* was undetectable in all reproductive sporophyte stages (Figure 3A). For those genes tested for gametophyte expression, expression was absent for *CericMADS26*/*CMADS1* and *CericMADS11* but detected in at least one stage for the other tested MIKC^c^ genes and for the MIKC* gene *CericMADS32* (Figure 3A; Supplemental Figure 1).

To gain further insights into putative functions of *Ceratopteris* MIKC^c^ genes, expression of *CericMADS26*/*CMADS1* and its two most closely related genes, *CericMADS2*, *CericMADS29*, plus *CericMADS33* from the neighbouring clade was quantified in a range of gametophyte and sporophyte tissues using quantitative RT-PCR (Figure 3B). We showed that *CericMADS26* is significantly enriched in later-stage reproductive tissue compared to corresponding vegetative stages (p < 0.05) (Figure 3B). Reproductive sporophyte tissues also show a non-significant trend of increasing with maturity of the tissue, compared to gametophyte and vegetative sporophyte tissue where expression remains similar (Figure 3B), confirming and extending the data of Hasebe et al. (1998). In contrast, *CericMADS2* shows enriched expression in hermaphrodite gametophytes and reproductive tissue compared to vegetative tissue and male gametophytes (p < 0.05) (Figure 3B). *CericMADS29*, the closest relative of *CericMADS2* (Figure 3A), shows a similar declining pattern of expression to *CericMADS2* in gametophytes, with enrichment in 5-day hermaphrodite gametophytes compared to all other tissues (Figure 3B). *CericMADS33* is also significantly enriched in 5-day hermaphrodite gametophytes compared to all other tissues, (Figure 3B). We thus found that expression patterns of MIKC^C^ genes in *Ceratopteris* cannot be predicted based on phylogenetic similarity alone, with *CMADS1* showing a distinct reproductive sporophyte-preferential pattern even from its closest relatives, which show a trend towards hermaphrodite gametophyte-preferential patterns.

To extend the RT-PCR and qRT-PCR experiments, and to gain a family-level understanding of MADS-box expression across sporophyte shoot development, we interrogated recent transcriptome data generated from different developmental stages of *Ceratopteris* vegetative and reproductive fronds ((Plackett and Catoni 2025); Figure 3C; Supplemental tables 3 and 4). Transcripts were detected for all but one (*CericMADS3*) of the *CericMADS* MIKC^c^ genes and all 20 of the detected *CericMADS* MIKC^c^ genes are expressed in some sporophytic tissues (Figure 3C). Four genes, *CericMADS12*, *CericMADS17*, *CericMADS19* and *CericMADS24*, are noticeably enriched in all young (25d) vegetative tissues compared to other samples (Figure 3C). Conversely, three genes, *CericMADS7*, *CericMADS31* and *CericMADS33*, are reduced in all 25d tissues compared to 60d and 130d samples (Figure 3C). *CericMADS6*, *CericMADS10*, *CericMADS11*, *CericMADS13*, *CericMADS17, CericMADS23* and *CericMADS24* show overall reduced expression in reproductive tissue: amongst genes with relatively low reproductive expression *CericMADS13* and *CericMADS23* show an increase in mature reproductive fronds (Figure 3C). In contrast, *CericMADS21* and *CericMADS26* and to a lesser extent *CericMADS29*, are enriched in all stages of the 130d (reproductive) sporophyte (Figure 3C), supporting the qRT-PCR analysis for *CericMADS26* and *CericMADS29* (Figure 3B) and matching the genes’ phylogenetic clustering (Figure 3A). Two genes, *CericMADS2* and *CericMADS27*, show enrichment in young (apex and primordia) reproductive tissue before expression declines, with some evidence of a similar developmental pattern in 60d fronds (Figure 3C). *CericMADS2*, *CericMADS26* and *CericMADS29* have their lowest expression in 60d vegetative tissue, whereas *CericMADS33* shows a more evenly distributed expression pattern between 60d (vegetative) and 130d (reproductive) tissue (Figure 3C), again reflecting the qRT-PCR analysis (Figure 3B).

A number of MIKC^C^ genes show changes in expression more closely related to frond development than shoot reproductive phase. *CericMADS12*, *CericMADS19* and *CericMADS30* show enrichment in apical and/or primordia tissue in at least 2 out of three shoot ages (Fig. 3C). *CericMADS7* shows some enrichment in later stages of frond development (fiddlehead and expanding fronds) in both 60d vegetative tissue and 130d reproductive fronds Similarly, *CericMADS5* and *CericMADS23* show enrichment in the most mature stages of both vegetative and reproductive frond development (Figure 3C).

Taken together, our expression analyses highlight the presence of most tested *Ceratopteris* MIKC^c^ cDNAs in the gametophyte and in the majority of stages of sporophyte development. The data suggest possible diverse roles for MIKC^c^ genes across all tissues based on their detailed expression patterns. We also detected different expression patterns within the sporophyte that may reflect developmental functions: 4 genes show greatest enrichment in young (25d) vegetative tissue while a further 6 show more general reduced expression in reproductive tissues compared to vegetative. 3 genes show increasing enrichment as sporophyte fronds mature while a further 2 show decreasing expression (60d and 130d), implying roles in the process of frond development. 2 genes show increased amplitude in reproductive tissues, but with a common pattern in 60d and 130d of decreasing expression as frond development progresses. Only 3 genes are enriched throughout the reproductive sporophyte: one of these is *CericMADS26*/*CMADS1*, independently confirming previous northern blot data (Hasebe et al. 1998). We have thus successfully identified two further MIKC^C^ MADs-box genes (*CericMADS21* and *CericMADS29*) as candidates for interactors with *CericMADS26/CMADS1* in reproductive sporophyte development.

### Protein structures of the *Ceratopteris* MIKC^c^ family compared to their Arabidopsis relatives

Hasebe *et al*. (1998) postulated that *CericMADS26*/*CMADS1* was most similar to the AGAMOUS clade (AG, AGL1/SHATTERPROOF1 (SHP1), AGL5/SHP2 and AGL11/SEEDSTICK (STK) in *Arabidopsis*) due to the presence of additional amino acids N-terminal to the MADS DNA binding domain in these proteins, plus the reproductive-specific expression of *CericMADS26*/*CMADS1*. To explore this further, we performed sequence alignments with both the *Arabidopsis* and *Ceratopteris* MIKC^c^ proteins (Figure 4). As in *Arabidopsis*, the conserved M and K domains within the *Ceratopteris* proteins are easily identifiable (Figure 4) with less well conserved intervening and C-terminal regions. Overall, there is a slightly, but not significantly, lower degree of conservation (amino acid consistency scores) within the M and K domains of the *Ceratopteris* proteins compared to the M and K domains of the *Arabidopsis* proteins (Figure 4). The average *Arabidopsis* M domain consistency score is 7.64 compared to 6.97 for the *Ceratopteris* M domain (p=0.13 in a Mann Whitney test) while the *Arabidopsis* K domain consistency score is 4.96 compared to 4.19 for the *Ceratopteris* K domain (p=0.17 in a Mann Whitney test). Over half the *Ceratopteris* proteins were found to have extended domains N-terminal to the M domain, all of which are much longer than the N-terminal extensions present in *At*AG and three other *Arabidopsis* proteins (Figure 4).

**Figure 4.**
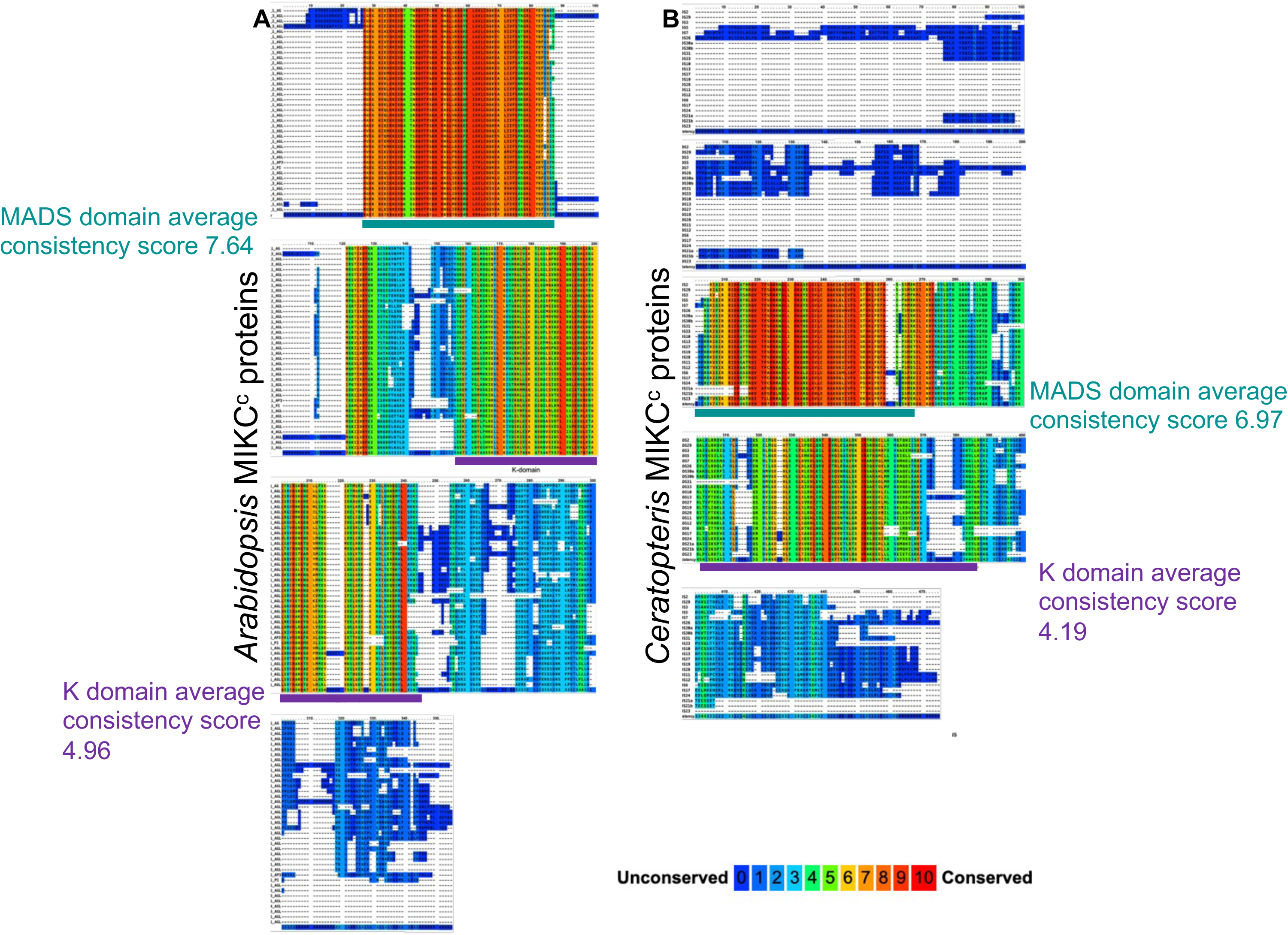
Comparison of *Arabidopsis* and *Ceratopteris* MIKC^c^ MADS-box protein sequence alignments. A) Graphical overview of *Arabidopsis* MIKC^c^ protein alignment. B) Graphical overview of *Ceratopteris* MIKC^c^ protein alignment. Alignments were generated using PRALINE (Simossis and Heringa 2005). Coloured shading represents the degree of conservation with red being the most conserved (a consistency score of 10 representing identical amino acids at a particular position) and dark blue being the least conserved (consistency score of 0). Consistency scores are shown as the continuous coloured line at the bottom of the aligned sequences: these were used to calculate average consistency scores of the M and K domains. Numbers above the sequences represent amino acid positions in the alignment. Below the alignment, the MADS domains are marked in teal and the K domains in purple.

We then modelled *Arabidopsis* and *Ceratopteris* MIKC^c^ protein structures from their sequences and related the predicted structures to the positions of the proteins on a phylogeny based on the sequence alignment (Figure 5; Supplemental Figure 2). Predicted *Arabidopsis* MIKC^c^ proteins fall into 2 groups: those whose overall structure resembles *At*AG and those that resemble FLOWERING LOCUS C (*At*FLC), with a sharply acute angle between the two alpha helices in the K domain (Supplemental Figure 2). A short, disordered region upstream of the M domain is present in AG, AGL1/SHP1 and AGL5/SHP2 (Supplemental Figure 2), which is not predicted to be helical, nor as long as the *Ceric*MADS26/CMADS1 N-terminal region (Figure 4; Supplemental Figure 2). 12 *Ceratopteris* MADS-box proteins in addition to *Ceric*MADS26/CMADS1 have highly extended regions N-terminal to the MADS-box (i.e. *Ceric*MADS2, *Ceric*MADS3, *Ceric*MADS5, *Ceric*MADS7, *Ceric*MADS21a/b, *Ceric*MADS23, *Ceric*MADS29, *Ceric*MADS30a/b, *Ceric*MADS31 and *Ceric*MADS33), with all but *Ceric*MADS23 grouped in a single clade despite their predicted structural differences (Figure 5; Supplemental Figure 2). More generally, the predicted structures of *Ceratopteris* MADS-box proteins show more variability than those in *Arabidopsis*, particularly in the I and K domains (Supplemental Figure 2). The *Ceratopteris* MADS-box protein structures that model most similarly to the *Arabidopsis* floral patterning proteins within their K domains are *Ceric*MADS2, *Ceric*MADS3, *Ceric*MADS5, *Ceric*MADS7, *Ceric*MADS10, *Ceric*MADS13, *Ceric*MADS17, *Ceric*MADS19, *Ceric*MADS26/CMADS1*, Ceric*MADS27, *Ceric*MADS28, *Ceric*MADS30, *Ceric*MADS31, *Ceric*MADS33 (Supplemental Figure 2). In contrast, the predicted structures of *Ceric*MADS11, *Ceric*MADS12, *Ceric*MADS21 and *Ceric*MADS23 more closely resemble *Arabidopsis* flowering time regulators, with a sharply acute angle between the two alpha helices of the K domain (Figure 5; Supplemental Figure 2). However, *Ceric*MADS11, *Ceric*MADS12, *Ceric*MADS21 and *Ceric*MADS23 are phylogenetically distant from the FLC flowering time clade (Figure 5).

**Figure 5.**
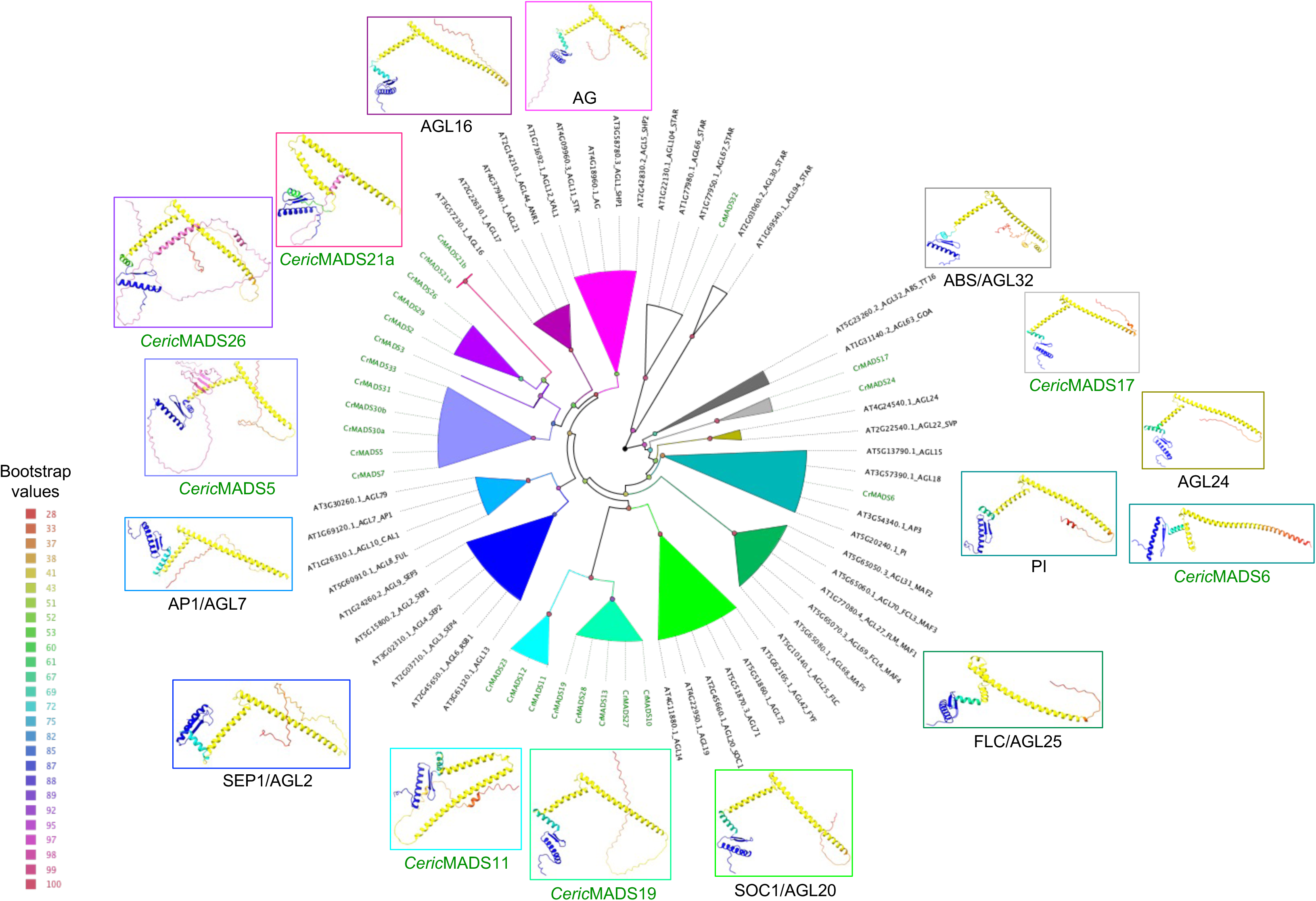
*Ceratopteris* MADS modelled protein structures show diversity compared to *Arabidopsis* predicted MADS protein models. All the *Ceratopteris* and *Arabidopsis* MIKC^c^ protein sequences plus representative MIKC* sequences as an outgroup were aligned and a phylogenetic tree generated. MIKC^c^ protein clades are shown in colour and for each clade a representative structural model is shown outlined in the same colour. *Arabidopsis* proteins are labelled in black text, and *Ceratopteris* proteins in green text. In the structural models, the M domain is shown in blue, the I domain in cyan/green and the K domain in yellow with the N-terminal extensions in magenta. The full complement of modelled structures is shown in Supplemental Figure 2. Bootstrap values of the nodes are shown in colour according to the legend on the left.

## Discussion

*Ceratopteris richardii* possesses 21 MIKC^c^ MADS-box genes. Expression of the majority is detected in sporophyte tissues, including reproductive tissue, with some also expressed in gametophyte tissue. Our data is in alignment with previous northern blot and *in situ* hybridisation analysis of *CericMADS26*/*CMADS1*, *CericMADS28*/*CMADS2*, *CericMADS13*/*CMADS3*/*CRM1* (Hasebe et al. 1998), *CericMADS17*/*CMADS6*/*CRM3* (Schulz and Theissen 2025) and RT-PCR analysis of the *CMADS3*-related *Dryopteris fragrans Df*MADS1 (Huang et al. 2014).

In contrast to previously identified reproductive sporophyte-expressed genes (Hasebe et al. 1998) (Schulz and Theissen 2025) (Huang et al. 2014), *CericMADS11*/*CMADS4* was not detected by northern blotting in gametophytic or sporophyte reproductive tissue (Hasebe et al. 1998). However, we were able to amplify *CericMADS11* from all stages of sporophyte reproductive development but not gametophyte tissue (Figure 3; Supplemental Figure 1). Moreover, *LAMB2* (the *Lycopodium* homologue of *CericMADS7*) is only weakly detected by semi-quantitative RT-PCR in reproductive strobili compared to vegetative shoots (Svensson and Engstrom 2002) but there was sufficient expression in *Ceratopteris* to amplify *CericMADS7* from expanding and mature reproductive frond cDNA, but not the earlier fiddlehead stage (Figure 3A; Supplemental Figure 1), and comparable expression in vegetative and reproductive tissues (Figure 3C), suggesting possible functional diversification of this gene between *Lycopodium* and *Ceratopteris*.

Our global analysis of *Ceratopteris* MIKC^c^ gene expression (Figure 3C) using a recent transcriptomic atlas of sporophyte shoot development (Plackett and Catoni 2025) revealed relatively little tissue- and/or stage-specific enrichment, with some differences between young (25d) and older (60d, 130d) sporophytes and some similar patterns found across frond development between vegetative and reproductive sporophytes. In contrast to the restricted expression of the *Arabidopsis* floral homeotic MIKC^c^ genes, the majority of *Ceratopteris* MIKC^c^ genes are expressed in the vegetative sporophyte. 18 out of 20 MIKC^c^ genes (excluding *CericMADS3*) were enriched in at least one sporophyte vegetative sample, with only 2 genes (*CericMADS21* and *CericMADS26*/*CMADS1*) specifically enriched in reproductive (130d) sporophytes. 4 genes (*CericMADS12*, - *17*, *-19* and - *24*) were enriched specifically in 25d young sporophyte tissue (Figure 3C). In a recent transcriptome analysis of the fern *Lygodium japonicum*, the majority of MIKC^c^ genes were also expressed in vegetative sporophyte tissues, with some enrichment towards either younger or older sporophyte tissues; however, reproductive sporophytes were not tested (Zhang et al. 2024). As with *CericMADS26*/*CMADS1*, some of the *Lygodium CRM6* clade show reduced or no gametophyte expression, although the closest *Lygodium* relatives to *CericMADS26* generally do not show reduced gametophyte expression (Zhang et al. 2024).This broad expression of fern MIKC^c^ genes contrasts with the specific expression patterns and reproductive developmental function of the *Arabidopsis* floral homeotic MIKC^c^ genes

Despite changes in the dominance of the sporophyte during plant evolution, the expression of MIKC^c^ genes in reproductive tissues appears to be ancient and conserved, as the single MIKC^c^ gene in the multicellular alga *Chara globularis* (stonewort) is also expressed in reproductive tissue and the unicellular desmid *Closterium CpMADS1* is expressed during induction of gametangia (Tanabe et al. 2005). In the moss *Physcomitrium*, expression of the six MIKC^c^ genes is widespread, and reproductive functions have been proposed (Quodt et al. 2007; Singer et al. 2007), but only certain functions are conserved in flowering plants (Koshimizu et al. 2018).

Our transcriptome analysis (Figure 3C) detected the first transcripts identified for *CericMADS5*, *CericMADS21a*/*b* and *CericMADS30a*/*b* - these transcripts were previously predicted, but not detected in the most recent genome annotation (Marchant et al. 2022). This leaves only *CericMADS3* as a potential *Ceratopteris* MIKC^c^ ‘gene without a transcript’.

Modelling of *Ceratopteris* MIKC^c^ proteins compared to their *Arabidopsis* counterparts demonstrated that predicted *Ceratopteris* protein structures are more diverse than those in *Arabidopsis*. This is particularly noticeable within the K domains, suggesting that there may be differences in dimerization properties of MADS-box proteins between the two lineages. The *Ceratopteris* proteins with a sharp acute angled bend in the K domain (*Ceric*MADS11, *Ceric*MADS12, *Ceric*MADS21 and *Ceric*MADS23) that resemble *Arabidopsis* FLC and other flowering time regulators lie in clades distinct from FLC and its relatives. Over half (13/23) the predicted *Ceratopteris* MIKC^c^ protein structures have extended N-terminal domains in contrast to less than 10% (3/39) of the *Arabidopsis* proteins. Whilst *Arabidopsis* AG, AGL1 and AGL5 N-termini have no specific predicted structure, the *Ceratopteris* proteins’ N-termini are mostly predicted to contain alpha helical regions, with one beta-sheet (*Ceric*MADS5).

*Ceric*MADS26/CMADS1 is predicted to have an extended N-terminus upstream of the M domain (Figure 4; Supplemental Figure 3). Although this upstream sequence was postulated to render CMADS1 more similar to AG and its relatives (Hasebe et al 1998), our modelling shows that the *Ceric*MADS26/CMADS1 N-terminal domain is longer and predicted to be alpha-helical, unlike AG, AGL1 and AGL5. However, the K-domain of *Ceric*MADS26/CMADS1 models similarly to AG. The original analysis (Hasebe et al., 1998) used a limited range of fern protein sequences and only 17 *Arabidopsis* sequences. Moreover, the extended N-terminus is missing in AG orthologues from gymnosperms, gnetophytes and Ginkgo (Jager et al. 2003) and the N-terminal extension evolved convergently in the *euAG* and *PLE* subclades of the AG lineage in dicot flowering plants (Ma et al. 2020). The function of the AG N-terminus is unknown, as it is not essential for DNA binding or formation of ectopic reproductive organs (Mizukami et al. 1996). Our data suggest that very long, structured N-termini may be a more general feature of fern MIKC^c^ MADS proteins, rather than AG-like MADS proteins, due to the proportion and diversity of *Ceratopteris* proteins containing the extended N-terminal domain. In addition, the *Lygodium* CRM6 clade (related to *Ceric*MADS33) also possess long N-terminal regions (Zhang et al., 2024b). Whether *Ceratopteris* N-terminal extensions have evolved convergently or from a common ancestor is unknown.

Taken together, our study delivers the largest number of cloned, confirmed *Ceratopteris* MIKC^c^ MADS-box cDNA sequences to date, provides new information on *Ceratopteris* MIKC^c^ gene sequences, and provides new expression data for all but one of the *Ceratopteris* MIKC^c^ genes. The predicted structures of the *Ceratopteris* MIKC^c^ proteins provide new insights into MIKC^c^ protein diversity and evolution. We speculate that the diversity of *Ceratopteris* MIKC^c^ structures and extended N-termini could underpin novel DNA binding functions or protein-protein interactions: these can be investigated in future work.

## Materials and methods

### Plant material and growth conditions

Wild type *Ceratopteris richardii* Hn-n (https://c-fern.org/) was used. *Ceratopteris* spores were cleaned by incubation with 1ml of 20% bleach (sodium hypochlorite) with 0.1% Tween20 for 15 minutes at room temperature with gentle mixing. Spores were washed 4 times in sterile distilled water and resuspended in 1ml sterile distilled water for imbibition via incubation at room temperature in darkness for 48-72h. To generate gametophytes, spores were plated on sterile 1x C-fern medium with 1% agar pH6.0 as described previously (Plackett et al. 2015) in 90mm petri dishes, using cellophane disc overlays (AA Packaging, UK) sterilised between filter paper discs (Whatman, UK). To ensure the plant growth environment stayed humid, Sterile distilled water was added to the plates on alternate days. To generate sporophytes, plants were grown on soil (Levington Peat Free All-purpose compost) supplemented with liquid 1x C-fern medium. The tissue for RNA extraction was harvested from immature and mature gametophytes (5, 6, 10, and 13 days old) and from the fiddleheads, expanding fronds and mature fronds of both vegetative (60 day) and reproductive (130 day) sporophytes.

### Generation of RNA and cDNA

Tissue was harvested and snap frozen in liquid nitrogen. RNA was generated using an RNeasy Plant Mini kit (Qiagen, Manchester UK) according to the manufacturer’s instructions. cDNA was synthesised using SuperScript II (Thermo Fisher Scientific) according to the manufacturer’s protocol.

### Identification, cloning and sequencing of *CericMADS* genes

Primers (Supplemental Table 2) were designed to PCR-amplify the longest predicted coding sequence of each *CericMADS* gene using the previous literature including the annotated *Ceratopteris* genome that was available at the start of the project (Marchant et al. 2019; Munster et al. 1997). PCR amplification was attempted from the cDNA of the different stages of *Ceratopteris* development using Phusion DNA polymerase (Thermo Fisher Scientific). Initial denaturation took place at 98°C for 30s, with 29 cycles of 10s at 98°C, 20s of annealing at a temperature 3°C above the predicted T_m_ of each primer pair (around 55°C) and extension at 72°C for 0s per kb of predicted sequence length and extensions. A final extension step was added for 6 minutes at 72°C.

Amplified cDNA sequences were cloned into a pJET blunt cloning vector (Thermo Fisher Scientific), sequenced (Source Bioscience, Cambridge, UK) and checked by appropriate restriction digestion (New England Biolabs). The sequences of the cloned cDNAs were cross-checked against the updated *Ceratopteris* genome (Goodstein et al. 2012; Marchant et al. 2022) once it became available.

### Quantitative RT-PCR (qRT-PCR)

qRT-PCR was performed using Brilliant III ultra-fast SYBR® Green low ROX qPCR master mix (Cat. 600892, Agilent) in 384 well plates on an AriaMx machine (Agilent). Each 5 µL reaction contained 2.5 µL Sybr Green Supermix, 0.25 µL of the appropriate forward primer at 10 µM, 0.25 µL of the appropriate reverse primer at 10 µM and 2 µL of the appropriate cDNA at 5 ng/µL. For each sample, 4 technical repeats were performed. The cycling parameters were as follows: initial denaturation at 95 °C for 10 minutes, followed by 40 amplification cycles of a 15 second 95 °C denaturation step plus a 20 second 60 °C annealing step. The short amplicon length means that elongation occurs during the cycle up to the denaturation temperature. A melt curve was produced via 15s at 95 °C, 60s at 60 °C 15 s at 95 °C with a ramp rate of 0.075 °C per second. The normalised fold change was calculated via the 2-ΔΔCT method (Livak and Schmittgen, 2001). CT values were normalised against 3 reference genes, namely *ACTINB*, a putative *UBIQUITIN* gene, and a putative TATA gene ((Ganger et al. 2015); Supplemental Table 2).

### Expression analysis of *CericMADS* genes from RNAseq data

The expression of *CericMADS* genes in different *Ceratopteris* tissues was analysed using available RNAseq data ((Plackett and Catoni 2025); Supplemental Table 3). Raw reads were trimmed and filtered with TrimGalore (v 0.6.10) and assessed with FastQC (v0.11.9), then aligned to the *C. richardii v2.1* reference genome (Marchant et al., 2022) using HiSAT2 (v2.2.1) with default parameters. Alignments were sorted and indexed with SAMtools (v1.10), and gene-level counts obtained with HTSeq (v0.13.5). Counts were processed in R (v4.5.1): transcripts representing the same MADS box gene were merged, low-count genes (fewer than 10 reads in at least 3 samples) were filtered, and variance-stabilising transformation (VST, DESeq2 v1.50.2) applied to the full dataset. MADS-box genes were extracted, row-scaled to Z-scores (Supplemental Table 4), and visualised as a heatmap with hierarchical clustering of genes (Euclidean distance, complete linkage) using ComplexHeatmap (v2.26.0).

### Protein alignment, phylogenetic analysis and structure prediction

*Ceric*MADS predicted peptide sequences generated during this work and *Arabidopsis* MADS protein sequences retrieved from the *Arabidopsis* TAIR 10 genome annotation at Phytozome ((Goodstein et al. 2012); https://phytozome.net/) were used for alignment and phylogenetic analysis. For alignment visualisation in Figure 4, sequences were aligned using PRALINE (Simossis and Heringa 2005). For phylogenetic analysis (Figure 3 and 5), sequences were aligned with MAFFT (Madeira et al. 2024) using automatic settings. A consensus phylogeny was constructed with IQ-TREE (Trifinopoulos et al. 2016) using default settings. Phylogenetic trees were visualised using FigTree (https://tree.bio.ed.ac.uk/software/figtree/). Protein structures were predicted with AlphaFold using Google Colab and ColabFold (Mirdita et al., 2022) and visualised using ChimeraX (Meng et al., 2023) using the predicted sequence motifs to colour the M and K domains.

## Author contributions

Conceived research: AP and DC. Performed experiments: DC, AP, KM and JC. Analysed data: DC, KM, AP and JC. Generated data figures: DC, KM, JC. Wrote paper: DC and JC. Reviewed and edited paper: KM and AP.

## Acknowledgements

The research was funded by a University of Birmingham doctoral scholarship (DC), UKRI-BBSRC doctoral scholarship BB/T00746X/1 (KM), Royal Society University Research Fellowship URF\R1\191326 (AP) and Leverhulme Trust Research Fellowship RF-2024-335 (JC). We thank the University of Birmingham UoB genomics facility for sequencing services.

