## Supplemental Figures 1 and 2 for "Analysis of MIKC^c^-type MADS-box genes and proteins in the fern *Ceratopteris richardii*"

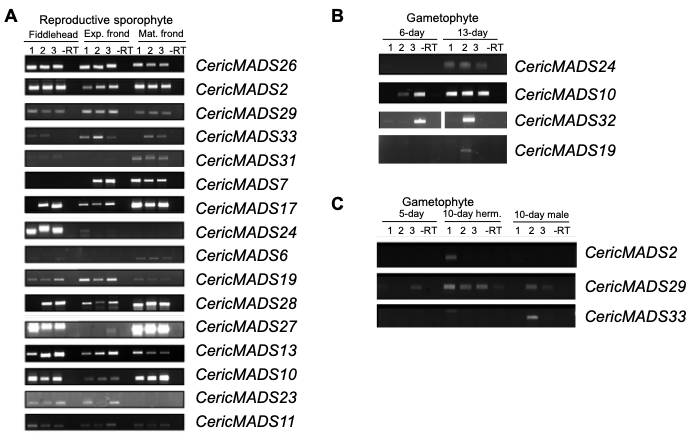

**Supplemental Figure 1. Detected RT-PCR amplification of *CericMADS* cDNAs from RNA extracted from gametophyte and reproductive tissues, for subsequent cloning and sequencing.**

A. Detected expression of the 16 *Ceratopteris* MIKC^c^ genes amplified by RT-PCR in reproductive sporophyte tissue from fiddleheads, expanding fronds (Exp.) and mature fronds (Mat.). For each gene and tissue type, 3 independent biological repeats from different plants are shown alongside a negative control (-RT).

B. Detected expression of the MIKC^c^ genes *CericMADS10, 19, 34* and *32* in 6- and 13-day old (immature and mature) gametophyte tissues. RT-PCR amplification from 3 tissue replicates are shown, alongside a negative control (-RT).

C. Detected expression of the MIKC^c^ genes *CericMADS2*, *CericMADS29* and *CericMADS33* in gametophyte tissue from immature (5 day) and mature (10 day) hermaphrodites and males. RT-PCR amplification from 3 tissue replicates are shown, alongside a negative control (-RT).

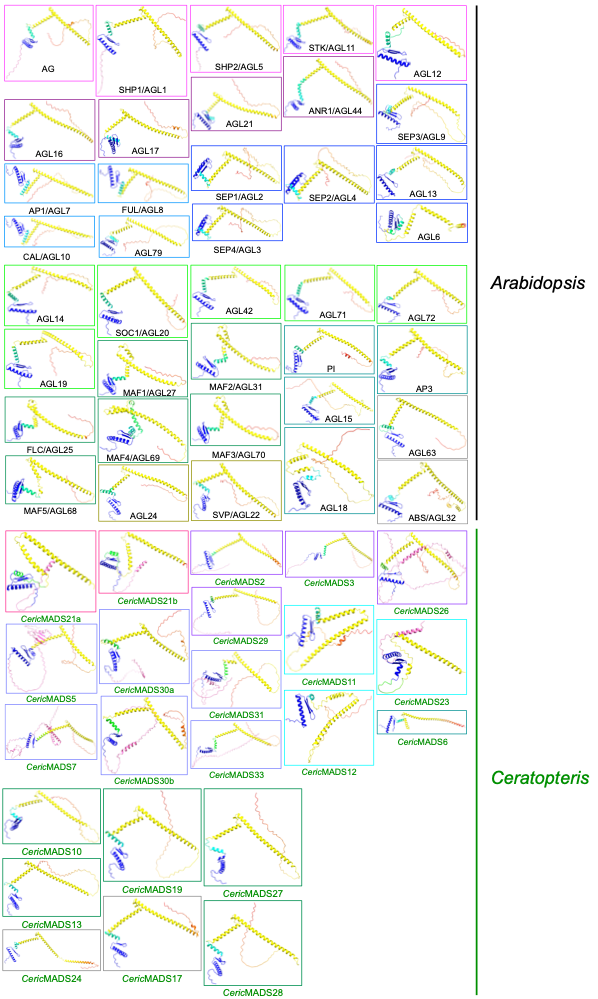

**Supplemental Figure 2. Modelling of *Arabidopsis* and *Ceratopteris* MIKC^c^ protein structures.**

*Arabidopsis* and *Ceratopteris* MIKC^c^ protein models. In the structural models, the M domain is shown in blue, the I domain in cyan/green and the K domain in yellow with the N-terminal extensions in magenta. Outline colours correspond to the phylogenetic groups in Figure 5.
